# Adoption of MMPose, a general purpose pose estimation library, for animal tracking

**DOI:** 10.64898/2026.03.29.715167

**Authors:** Jessica D. Choi, Vivek Kumar

## Abstract

Markerless pose estimation has emerged as a powerful technique for animal behavior quantification, capable of high resolution tracking of body parts. Many neuroscience labs rely on tools like DeepLabCut and SLEAP, which provide accessible interfaces but restrict users to a narrow set of models and configurations. In this work, we adopt MMPose an open source, general-purpose computer vision library to build a workflow for training and evaluating multiple state-of-the-art models on animal video datasets. We benchmark these models in two scenarios: (1) a complex maze assay with occlusions and varied backgrounds, and (2) a simpler open field arena with a high-contrast background. Our results show that a bottomup model (DEKR) delivers the highest accuracy in the complex task, whereas lighter-weight models (e.g., SLEAP) offer superior speed highlighting a clear trade-off between accuracy and throughput. We also evaluate a recently published foundation model (TopViewMouse-5K) trained on a large top-view mouse dataset to test its generalization. It performs poorly on our tasks at zero-shot, and even when we combine its data with our training set, we observe no consistent benefit. These findings emphasize the importance of context-specific model selection and the need for more diverse training data to create generalizable pose models. By leveraging a general-purpose vision library, researchers can flexibly choose models that best suit their experimental needs. This work illustrates how adopting advanced computer vision frameworks can accelerate behavioral neuroscience and genetics research, paving the way for more scalable, reproducible, and sensitive analysis of animal behavior.

## 2 Introduction

Neuroscientists seek to link genetics to altered neuronal phenotypes and ultimately altered behavior, with the goals of understanding mechanisms of disease and therapeutic development. Animal models represent an indispensable tool in pursuit of this goal, with their behavioral phenotypes representing critical disease relevant behavioral biomarkers. Historically, animal behavior has been analyzed through direct observation or manual annotation of video recordings. However, manual annotation approaches are labor intensive, subjective, and capture only limited snapshots of an animal’s behavioral repertoire, all of which ultimately limit the scalability and objectivity required for robust behavioral studies [1, 2]. These methods can also distort our understanding of disease progression by confounding behavioral observations with the stress of handling and non-ethological environments [3, 4, 5, 6, 7].

To overcome many of these limitations, researchers have implemented automated methods from advances in machine learning and computer vision [8, 9]. In early work, researchers relied on background subtraction, but this method limited the behavioral diversity that could be considered and required simplified visual environments to achieve good performance, which limited extracting ethological relevance to disease [10, 11, 12]. More recent technical advances in computing have enabled the use of markerless pose estimation as a first step in tracking animal body parts. Pose estimation allows for tracking at high spatiotemporal resolution and enables researchers to quantify behavior without markers or invasive techniques. Pose estimation is a useful data-efficient approach compared to a classic CNN which can save much time investment in downstream behavior analysis. However, the impact of these advances has been limited in rodent research as many biology-focused labs find it difficult to adopt state-of-the-art pose estimation methods. Tools like DeepLabCut and SLEAP lower technical barriers by offering graphical interfaces and streamlined workflows, however they often abstract away model customization and configuration choices [13, 14]. This limits flexibility and prevents researchers from adapting models to the specific needs of their behavioral assays or imaging conditions, particularly as behavior paradigms become more complex.

In contrast to the models currently used in the rodent field, human pose researchers have developed a broad collection of models, some optimized for speed, others for accuracy, each showing different strengths depending on the context [15, 16, 17]. Yet in animal tracking studies, most rely on a single model, without systematic comparison. This can introduce hidden model biases that can limit biological interpretation.

Beyond model flexibility, researchers lack standardized dataset formats for rodent pose estimation. Each tools has its own specialized data format that locks users into a specific set of downstream tools. As a result, data sharing remains rare, and benchmarking across models and datasets proves difficult. This further limits implementation and adoption of configurable algorithms from the broader computer vision field that may offer more specialized outcomes for specific research problems.

To address these challenges, we test MMPose, a general-purpose human pose estimation library developed by the OpenMMLab project, for use in mouse tracking tasks [18]. MMPose supports a variety of state-of-the-art models, integrates easily with object detection tools like MMDetection [19], and uses the MS COCO keypoint format to enable annotation standardization [20]. This platform allows users to configure model architectures, data augmentations, and training regimes with flexibility and transparency. This is valuable as researchers can then optimize and use the best models for their own specific use-cases in a data structure that allows for sharing of annotations.

We adopt MMPose for rodent pose estimation and evaluate its performance in tasks relevant to neuroscience and genetics. We compare a variety of models in two scenarios: a complex maze environment and a simple open field arena. This is to understand the performance attainable with different types of datasets, also depending on the specific questions a researcher is striving to answer. We assess each models pose estimation accuracy and inference speed to better understand the trade-offs in performance when selecting models. We also examine whether a recently proposed foundation model trained on a large top-view rodent dataset can generalize to new settings, and how performance can improve with different model architectures [21]. Our results show the value of modular, configurable pipeline selection and underscore the need for diverse, shared datasets in behavioral science.

## 3 Results

### 3.1 Pose Estimation in a Complex Maze

We evaluated the full MMPose-based pipeline on the Kumar lab maze dataset (Figure 1A) which we used to test how pose estimation models handle occlusion and visually complex backgrounds. The maze dataset comes from an in-house assay adapted from the Meister lab maze [22] that measures navigation strategy and spatial learning. Because this task requires precise tracking of the mouse’s body position in the branches of the maze, accuracy is a valuable metric. We quantified model accuracy using the Proportion of Correct Keypoints (PCK) metric across body length thresholds. Nine models were benchmarked: Def-DETR HRNet, Def-DETR DeepPose, RetinaNet HRNet, RetinaNet DeepPose, YOLO HRNet, YOLO DeepPose, DEKR, as well as DeepLabCut (DLC), and SLEAP.

**Figure 1:**
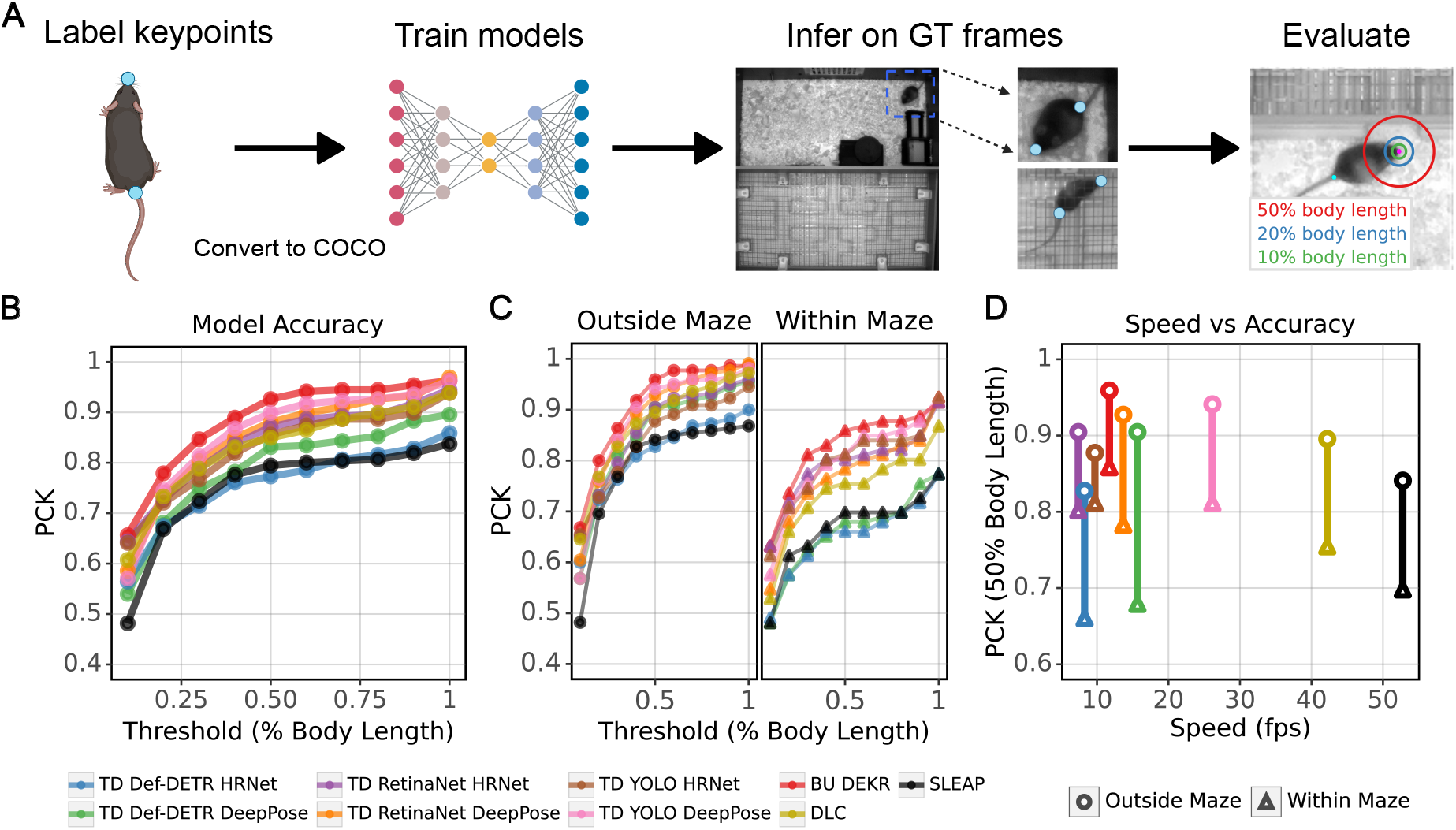
Tradeoffs between accuracy and throughput across model architectures. (A) Keypoint tracking pipeline using MMPose. (B) Model accuracy by PCK across body length thresholds. (C) Speed vs. accuracy at a body length threshold of 0.5. (D) Accuracy by location in arena.

Model performance can vary depending on environmental complexity and architecture choice. Across all models, PCK values exceeded 75% at the 0.5 body length threshold, showing that the task was learnable with the training data (Figure 1B). Particularly, the bottom-up DEKR model consistently achieved the highest accuracy across all thresholds, exceeding 90% PCK at 0.5 body length. This result shows that a bottom-up architecture can be more robust in this cluttered and complex environment with consistent camera angle and listing.

Neural network tracking models have differing performance based on complexity of the task and occlusions [23]. Many experimentalists report one number as the final performance metrics, however this can be misleading. For instance, in a previous paper we found that keypoint tracking errors have small but significant performance differences based on the coat color of the mouse [Sheppard et. al., [24] Fig. 2]. Similarly, environmental complexity has been shown to alter performance. Our arena is highly polarized with a much simpler task in the open area and high level of occlusion within the maze. Therefore, we quantified performance both inside and outside of the maze.

**Figure 2:**
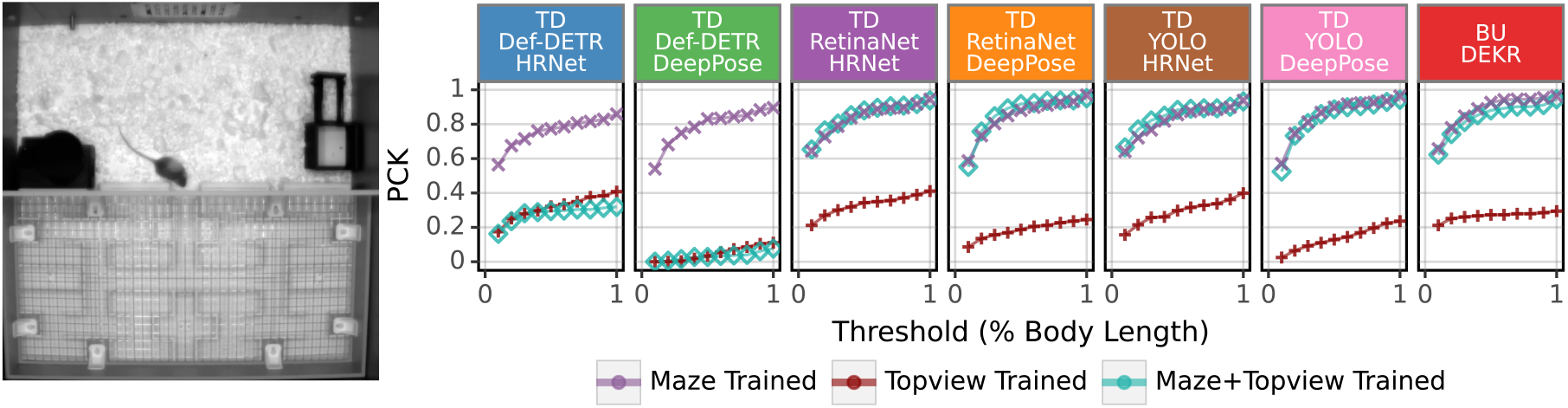
TopViewMouse-5K dataset fails to generalize using zero-shot inference on complex maze-task. Models trained on three datasets: maze-only, TopViewMouse-5K-only, and maze combined with TopViewMouse-5K. All models were benchmarked and show PCK performance on the maze ground truth. Example image of the maze dataset shown.

To quantify this, we manually annotated each frame as “inside” or “outside” the maze and calculated the PCK at the 0.5 body length threshold for each region (Figure 1C). We then computed the difference in performance between “inside” and “outside” tracking. Tracking performance varied drastically between the open and occluded (maze) regions of the image. The absolute differences per model ranged from 6.6% to 22.5%, confirming previous observations that a more complex environment confounds performance. However, we find that the drop in performance is model-specific. DEKR still led in accuracy, dropping only 10.1%. The accuracy outside of the maze reached 0.96. In contrast, we find that DLC and SLEAP have a greater reduction of absolute performance within the maze, with PCK losses of 14.1% and 14.3% at the same threshold, respectively. The top-down Def-DETR models showed the largest losses (16.7% and 22.5%). YOLO HRNet was the most stable overall, with an absolute loss of only 6.6% between inside and outside regions. These results emphasize that robustness to occlusion and visual complexity is architecture-dependent, which emphasizes the importance of selecting the correct context-specific pose estimator for the task at hand.

In high-throughput experiments such as pharmaceutical or behavioral genetics screens, inference speed becomes a crucial component of model evaluation. We measured how many frames per second (FPS) are processed during inference. SLEAP achieved the highest throughput at 52.8 FPS, followed by DLC (42.2 FPS, Figure 1D). YOLO DeepPose balanced speed and accuracy, with a speed of 26.1 FPS, and a PCK of 0.90 at 0.5 body length. In contrast, RetinaNet HRNet was the slowest (7.39 FPS). The other models clustered from 7 to 16 FPS. Although DEKR operated at 11.7 FPS, it maintained the highest accuracy, demonstrating the clear trade-offs between throughput and precision. In practice, researchers should therefore choose models based on whether speed or accuracy is more limiting for their application.

### 3.2 Performance on Complex Maze Task

Foundation models that promise “out-of-the-box” tracking could alleviate the burden of annotating and training specialized models per experimental setup. To assess the performance of the recently proposed foundation model TopViewMouse-5K [21], we trained MMPose models using the TopViewMouse-5K training data and evaluated them on the maze dataset to assess the ability to generalize to other environments.

All TopViewMouse-5K-trained models performed poorly, with PCK values between 0.03 and 0.34 at 0.5 body length threshold (Figure 2 and Table 1). This indicates that the environment and visual shift from the tasks included in TopViewMouse-5K to the complex and occluded maze is too large for the models to generalize without adaptation.

**Table 1:** Performance of models trained with different datasets and dataset combinations on maze and open field test sets.

|  | Model | Dataset | PCK | Dataset | PCK |
| --- | --- | --- | --- | --- | --- |
|  | TD Def-DETR HRNet | Maze | **0.77** | OFA | **0.93** |
|  |  | Maze + TopViewMouse-5K | 0.29 | OFA + TopViewMouse-5K | 0.68 |
|  |  | TopViewMouse-5K | 0.32 | TopViewMouse-5K | 0.61 |
|  | TD Def-DETR DeepPose | Maze | **0.83** | OFA | **0.92** |
|  |  | Maze + TopViewMouse-5K | 0.03 | OFA + TopViewMouse-5K | 0.47 |
|  |  | TopViewMouse-5K | 0.03 | TopViewMouse-5K | 0.42 |
|  | TD RetinaNet HRNet | Maze | 0.87 | OFA | **0.96** |
|  |  | Maze + TopViewMouse-5K | **0.88** | OFA + TopViewMouse-5K | 0.95 |
|  |  | TopViewMouse-5K | 0.34 | TopViewMouse-5K | 0.67 |
|  | TD RetinaNet DeepPose | Maze | 0.88 | OFA | **0.95** |
|  |  | Maze + TopViewMouse-5K | **0.92** | OFA + TopViewMouse-5K | **0.95** |
|  |  | TopViewMouse-5K | 0.19 | TopViewMouse-5K | 0.71 |
|  | TD YOLO HRNet | Maze | 0.86 | OFA | 0.92 |
|  |  | Maze + TopViewMouse-5K | **0.88** | OFA + TopViewMouse-5K | **0.94** |
|  |  | TopViewMouse-5K | 0.30 | TopViewMouse-5K | 0.61 |
|  | TD YOLO DeepPose | Maze | **0.90** | OFA | 0.91 |
|  |  | Maze + TopViewMouse-5K | 0.89 | OFA + TopViewMouse-5K | **0.95** |
|  |  | TopViewMouse-5K | 0.13 | TopViewMouse-5K | 0.55 |
|  | BU DEKR | Maze | **0.93** | OFA | **0.96** |
|  |  | Maze + TopViewMouse-5K | 0.88 | OFA + TopViewMouse-5K | **0.96** |
|  |  | TopViewMouse-5K | 0.27 | TopViewMouse-5K | 0.39 |
|  | DLC | Maze | 0.85 | N/A | N/A |
|  | SLEAP | Maze | 0.79 | N/A | N/A |

We next asked if supplementing the maze dataset with TopViewMouse-5K data could improve performance through increased diversity. We retrained each model on the combined dataset (Maze + TopViewMouse-5K) and tested on maze ground truth. In most cases, performance of the combined dataset models stayed the same as maze-trained models or declined. The bottom-up DEKR model dropped from 0.93 to 0.88 PCK (0.5 body length). In three cases (top-down RetinaNet HRNet, RetinaNet DeepPose, YOLO HRNet), the models improved marginally <1% from the maze-trained models (0.28% - 0.9%). In contrast, the Def-DETR models dropped performance drastically to TopViewMouse-5K levels, suggesting that the transformer-based models likely requires much more training data to be able to generalize and benefit from mixed training.

These results show that the TopViewMouse-5K data alone cannot generalize to the maze task. When combining the datasets, it can even reduce accuracy. Therefore, the current foundation model datasets for mouse tracking lack the diversity required for robust generalization to complex environments.

### 3.3 Performance on Simple Open Field Task

To test the effect of visual complexity and generalizability of our findings, we repeated our benchmarks using the environmentally simpler Open Field Arena (OFA) dataset [25]. This dataset features a white background with minimal visual clutter. The OFA dataset includes 12 keypoints across the mouse’s body.

Mirroring our previous experiments, we trained models on three datasets: OFA only, TopViewMouse-5K only, and a combined dataset (OFA + TopViewMouse-5K). As expected, OFA-trained models consistently reached high PCK scores when evaluating against OFA ground truth data (Figure 3, Table 1). All models surpassed 0.90 PCK at a 0.5 body length threshold. This demonstrates that the simpler open field context has less variation in model architecture performance.

**Figure 3:**
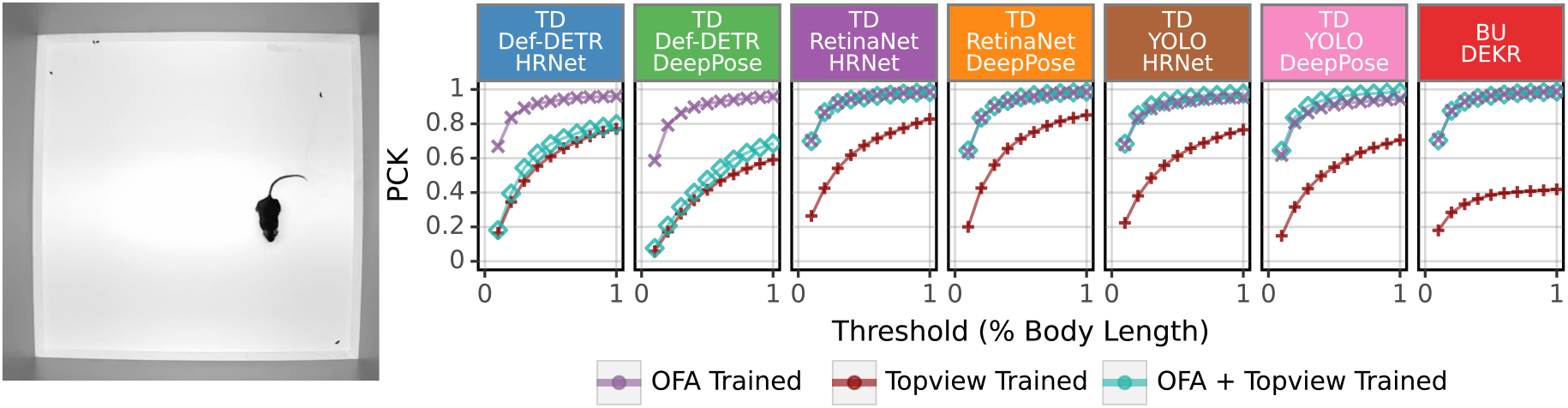
Models achieve high performance on simple open field arena. Models trained on three datasets: openfield-only, TopViewMouse-5K-only, and openfield combined with TopViewMouse-5K. All models were benchmarked and show PCK performance on the openfield ground truth. Example image of the openfield dataset shown.

When testing models trained only on TopViewMouse-5K against OFA ground truth, performance did not surpass 0.75 PCK. However, these models have much better performance on the OFA compared to the maze data. This supports the hypothesis that the TopViewMouse-5K models can generalize to visually simple environments but fail in complex ones.

We also test if adding TopViewMouse-5K data to the OFA training set boosts performance. Adding TopViewMouse-5K data to the OFA training set did not yield consistent improvement, and brought the Def-DETR model performances down. Together, these results support the conclusion of the current state of foundation-model data alone being insufficient for environmentally complex assays. There is an opportunity for future work to address and supplement mouse tracking foundation models.

## 4 Discussion

Markerless pose estimation has become prevalent in behavioral neuroscience, allowing researchers to quantify naturalistic mouse behavior at scale. By extracting pose from long-term video recordings, investigators can correlate behaviors with strain genetic backgrounds and disease phenotypes, and quantify behavioral modifications to therapeutic interventions. Traditional behavior phenotyping is limited by the high time-cost of manual annotations, which limits throughput, sensitivity, and reproducibility [1, 2, 26]. Neuroscientists have adopted pose estimation toolkits like DeepLabCut (DLC) [13] and SLEAP [14] which are easy to use and require little programming. Though this is valuable for democratization and accessibility, these tools lock users into fixed architectures, restricting the adoption of advances from the broader computer vision community. As behavioral assays become more complex through environment, multi-animal assays, long-term monitoring, or applicability across different lab setups, there is a demand for more flexible and powerful workflows.

In this study we tested whether MMPose, a general-purpose library [18] widely used in human contexts, could provide this flexibility in animal behavior research. MMPose equips researchers to find, adjust, and use the best model for their specific task based on their own experimental design and datasets. Different tasks may benefit from different model strengths. Our comparisons show that neuroscientists can improve analysis pipelines by exploring models beyond the typical defaults in current neuroscience toolkits. Importantly, our benchmarks show that no single model is optimal. Bottom-up DEKR maintained the highest accuracy on our complex maze dataset, but was one of the slower models for inference speed. Meanwhile SLEAP maintained the fastest inference speed, but was on the lower end for accuracy. This shows the trade-offs between throughput and precision that researchers should consider when designing experiments around pose and behavior.

Recent efforts from Li et al. [27] have shown the utility of applying MMPose to non-human domains, including *Caenorhabditis elegans, Drosophila*, and zebrafish. We extend this idea to rodent behavior, which comes with it’s own opportunities and challenges. Though some behavioral assays are short and contain clear goals, animal research has been moving towards computational ethology. The goal is to have as natural of an environment as possible, to capture spontaneous behavior in a way that is naturalistic, scalable, sensitive, and reproducible. This means one of the critical differences going from human to mouse tracking is the timescale of data being analyzed. Ethological mouse behavior analysis may involve continuous monitoring over hours and/or days. This requires a level of continuity across all frames to track rare events. This means that efficient long-term inference is valuable, not just accuracy in a snapshot. This contrasts from human benchmarks such as MPII, Human3.6M, COCO, and Posetrack that span seconds to minutes and analyze very specific actions such as walking or sitting [20, 28, 29, 15]. Scale needs to be taken into account and benchmarked in longterm naturalistic behavior analysis. Beyond scale, rodent datasets also includes challenging frames due to body deformability, occlusion, and variability across setups and labs. The ability to detect rare events is also important. Labels are expensive to produce and require expert knowledge, which makes model architecture decisions important to optimize tracking models. This means sample efficiency and transferability of these models matter.

Standardization can alleviate many of these challenges. Using a standard format like MS COCO would allow for sharing annotations and data across labs, allowing easier benchmarking and the ability for researchers to begin to use the tools that are constantly being developed and improved from the human field. Researchers can reproducibly benchmark and create fair comparisons across datasets and models. Notably, this also works to integrate with neuroscience data standards such as Neurodata Without Borders (NWB) [30] and data archives like DANDI, which are supported by the National Institute of Health (NIH) and the BRAIN Initiative. This shift is essential as it will allow for sharing annotations and neuroscience datasets. Other archives such as EMBER and Harvard Dataverse can also be used to publish behavior datasets and videos in standardized formats. These shared frameworks will push the community to create reproducible, collaborative studies, and will provide the scaffolding to create a large foundation model that can generalize across lab settings.

We also highlight the limitations of current foundation models. For example, while TopViewMouse-5K [21], is a valuable step towards standardized rodent pose data, currently lacks the diversity required to support generalizable rodent pose models. To build robust models capable of transferring across settings, the community needs to develop larger and more diverse datasets contributed from multiple labs. As more labs adopt these standards and share data, we can expect better generalization and more reliable foundation models for behavioral neuroscience.

Usability and adoption are nearly as important as performance. Many neuroscientists are rightly focused on experimental design, rather than software engineering. While computer science papers often release models or APIs, they do not release a UI or user-facing pipelines. Neuroscience would benefit from well-documented and exportable tools. Using a library like MMPose is a step in the right direction, but continued development should prioritize reproducibility and the ability for data sharing. We also encourage developers to continue to extend the utility of tools like DLC and SLEAP to continue to expand and include more model architectures from the human field. These toolkits are an extremely valuable entry point for many labs, and broadening the flexibility would increase their impact.

Our study provides guidance on model selection based on task complexity and resource constraints. Researchers should examine trade-offs in accuracy versus throughput depending on the complexity of their tracking task. Particularly for long-term ethological monitoring, inference speed may be the limiting factor. Computer vision researchers that are interested in improving throughput can use batching strategies and benchmark more efficient architectures. For more sensitive assays, accuracy may be the more important performance metric. We encourage researchers to use MMPose to evaluate models to find the ones best aligned with their scientific questions. Continued progress will depend on adopting shared frameworks, publishing diverse datasets, and promoting cross-disciplinary collaboration between computer vision and neuroscience.

## 5 Limitations of the study

Our study shows the utility of MMPose in rodent pose estimation, but some limitations should be considered when interpreting our findings.

First, our benchmarks are restricted to a top-view 2D single-animal recording setup. There are other behavioral paradigms involving multiple animals or alternate camera angles, and performance may differ in these contexts. However, our recommendation is not to prescribe one single model, rather we recommend users to benchmark and determine the best model for their particular use-case.

Second, our inference speed measurements were conducted on NVIDIA V100 GPUs. Researchers using different hardware will likely observe variation in throughput, and should interpret our speed benchmarks relative to their own computational resources.

Third, annotation quality is a potential source of variability in pose estimation datasets. Labeling strategies can differ across individuals and groups, which may affect reproducibility and cross-dataset comparisons. Adopting standardized annotation formats is a step towards centralizing, how annotations are stored and shared, though determining pose label quality remains a difficult endeavor.

## 6 Resource availability

### 6.1 Lead Contact

Requests for further information and resources should be directed to and will be fulfilled by the lead contact, Vivek Kumar.

### 6.2 Materials availability

This study did not generate new materials.

### 6.3 Data and code availability

The datasets used in analysis for this paper are all publicly available. The Kumar lab OFA dataset [25] can be accessed at: https://doi.org/10.5281/zenodo.6380163. The Kumar lab maze dataset has been deposited in Zenodo at: https://doi.org/10.5281/zenodo.19206232. The SuperAnimals TopViewMouse-5K dataset is publicly available and found through [21]. Pipeline and analysis code related to this manuscript is available on GitHub: https://github.com/KumarLabJax/mmpose-experiments.

## 7 Acknowledgments

This work was funded by The Jackson Laboratory Directors Innovation Fund, National Institutes of Health AG078530 (NIA, V.K.), DA041668 and DA048634 (NIDA, V.K.), MH138309 (NIMH, V.K.), and Nathan Shock Centers of Excellence in the Basic Biology of Aging AG38070 (NIA). T32 AG062409 (NIA, J.D.C.).

We thank Michelle Berger, Jacob Beierle, and Tuan Nguyen for reading over drafts of the manuscript and providing feedback. We thank Camille Berger-Liedtka for project coordination.

## 7.1 Author contributions

Contributions are described using the CRediT (Contributor Roles Taxonomy) categories.

- **Conceptualization:** J.D.C., V.K.
- **Investigation:** J.D.C.
- **Formal analysis:** J.D.C.
- **Visualization:** J.D.C., V.K.
- **Writing – original draft:** J.D.C.
- **Writing – review & editing:** J.D.C., V.K.
- **Funding acquisition:** V.K.

### 7.2 Declaration of interests

The authors declare no competing interests.

## 8 Materials and Methods

### 8.1 Maze Dataset Creation

#### 8.1.1 Sampling Strategy

To measure cognitive decline in mice through a long-term, naturalistic assay we adapted a complex maze task. The experiments were recorded for two days at 30 frames per second (FPS). To quantify these large-scale data, we created a pose estimation dataset that enables inference on a large volume of data that would be infeasible to hand-annotate.

To contribute to generalization of our trained models, we sampled frames for training to maximize diversity that included: (1) images with differing positions of the objects in the experimental setup, (2) camera angle variation across experiments, (3) images from more than 100 maze experiments, (4) images from all hours of the day, and (5) mice of various coat colors and genetic backgrounds. We initially annotated our keypoints using the DeepLabCut (DLC) toolkit, and used the built-in k-means method to cluster frames by visual similarity and enable diverse frame selection [13]. This method was applied on a per-video basis, so short clips may have limited visual diversity.

For ground truth sampling, we included mice of various coat colors from multiple experiments. Training and testing sets were separated on the experiment level, rather than by frame to ensure reliable inference on new experiments, providing value in scalability and consistency of the model being used in analysis.

#### 8.1.2 Labeling Process

We labeled 3,759 frames for training, validation, and testing. We labeled two keypoints per mouse: the nose and tail base. All annotations were labeled by one expert. As a quality check, we verified that the nose-tail base distance was within 200 pixels, confirming consistent labeling of the mouse within the 800×800 pixel images.

### 8.2 Datasets and Annotations

With the aim of robustly evaluating pose estimation models, we used three mouse pose datasets Table 2. We perform benchmarking on two datasets: the Kumar lab maze, and the simpler Kumar lab open field arena [25]. The third dataset, TopViewMouse-5K [21], was used solely for training. Models were trained on each dataset individually as well as on combinations of datasets to evaluate whether additional label diversity improves performance.

**Table 2:**
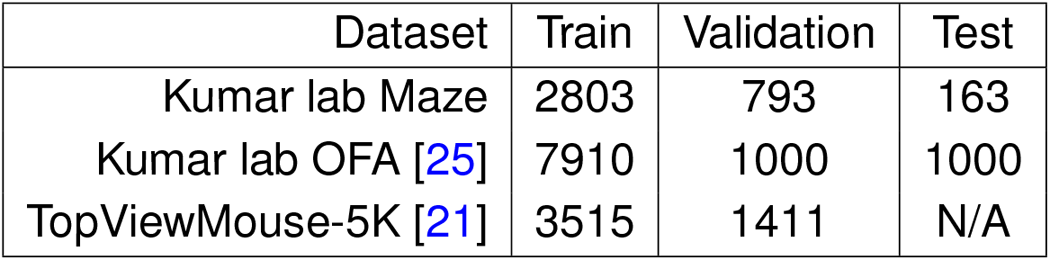
Dataset splits used for training and testing.

**Table 3:**
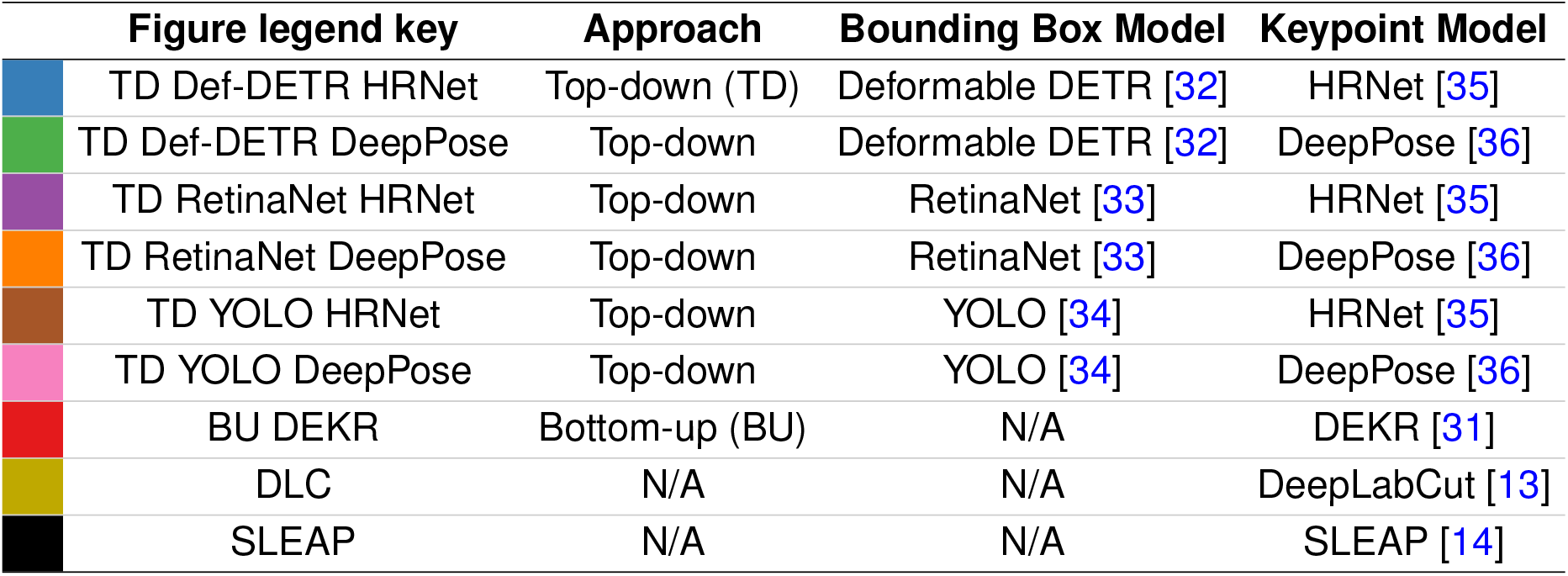
Models used for benchmark within and outside of the MMPose library.

|  | Figure legend key | Approach | Bounding Box Model | Keypoint Model |
| --- | --- | --- | --- | --- |
|  | TD Def-DETR HRNet | Top-down (TD) | Deformable DETR [32] | HRNet [35] |
|  | TD Def-DETR DeepPose | Top-down | Deformable DETR [32] | DeepPose [36] |
|  | TD RetinaNet HRNet | Top-down | RetinaNet [33] | HRNet [35] |
|  | TD RetinaNet DeepPose | Top-down | RetinaNet [33] | DeepPose [36] |
|  | TD YOLO HRNet | Top-down | YOLO [34] | HRNet [35] |
|  | TD YOLO DeepPose | Top-down | YOLO [34] | DeepPose [36] |
|  | BU DEKR | Bottom-up (BU) | N/A | DEKR [31] |
|  | DLC | N/A | N/A | DeepLabCut [13] |
|  | SLEAP | N/A | N/A | SLEAP [14] |

- **Kumar lab Maze (Maze)**. In-house dataset used to evaluate model robustness in a visually cluttered environment with occlusions. The behavioral assay measures cognitive flexibility and requires precise tracking of the mouse nose keypoint. The setup includes a home-cage area with bedding and a maze covered by a metal grate, increasing occlusion and thus increasing task complexity. Each frame was labeled with two keypoints (nose and tail base). The dataset was split into 2,803 training frames, 793 validation frames, and 163 test frames.
- **Kumar lab Open Field Arena (OFA)**. This dataset features a white background and a standard open field arena with genetically and visually diverse mice [25], including albino, nude, piebald, black, agouti, gray, and brown coat colors. Each frame was annotated with 12 keypoints. We used 7,910 frames for training, 1,000 for validation, and 1,000 for testing.
- **TopViewMouse-5K**. A publicly available dataset of approximately 5,000 annotated frames from eight behavioral setups [21]. These experiments were collected from multiple laboratory groups and include diverse conditions such as an open field, swimming tasks, and albino mice. This dataset was introduced as an example of a rodent foundation model that can generalize to mouse tracking tasks in a laboratory setting. We used 3,515 frames for training and 1,411 for validation. The original 27 keypoints from this dataset were mapped to the two keypoints used in the Kumar lab Maze, and the 12 keypoints used in the Kumar lab OFA Table 4.

**Table 4:** Keypoint mapping from the TopViewMouse-5K keypoint set to the OFA and Maze datasets. Keypoints only present in TopViewMouse-5K dataset not shown in table as they are not evaluated in our ground truth PCK metrics. Dashes indicate no counterpart.

| TopViewMouse-5K [21] | OFA [25] | Maze |
| --- | --- | --- |
| nose | nose | nose |
| left_ear | left_ear | - |
| right_ear | right_ear | - |
| neck | base_neck | - |
| mid_back | center_spine | - |
| tail_base | base_tail | base_tail |
| tail3 | mid_tail | - |
| tail5 | tip_tail | - |
| left_shoulder | left_front_paw | - |
| left_hip | rear_left_paw | - |
| right_shoulder | right_front_paw | - |
| right_hip | right_rear_paw | - |

### 8.3 Format Standardization

Pose estimation libraries, including MMPose, require training data to be in a specific annotation format that defines how images, keypoints, and all associated metadata are stored. However, when annotating in common neuroscience toolkits, such as DLC or SLEAP, the data becomes locked into these ad hoc formats that are not directly compatible. Further work is being developed within these toolkits to export annotations to common formats. In this paper, we converted all annotations originally created in SLEAP to the MS COCO format [20]. This conversion script is written in Python, and is available on the KumarLabJax/mmpose-experiments GitHub.

### 8.4 Models and Training

The specific pose estimation models we selected for this benchmark are popular baseline tools and models available through MMPose/MMDetection. The top-down pipelines in this benchmark were formed by pairing an object detector with a pose estimation head. Specifically, we used three detectors (Deformable DETR, RetinaNet, YOLOv3), and two pose heads (HRNet, DeepPose). These were benchmarked alongside a bottom-up model (DEKR) and two commonly used toolkits in neuroscience (DLC, SLEAP).

- **DEKR[31]**. Bottom-up keypoint detector that predicts heatmaps and associative embeddings to group joints into individuals. Bottom-up is typically more efficient than top-down methods in terms of training.
- **Deformable DETR (Def-DETR) [32]**. A transformer-based object detector that uses deformable attention to improve efficiency and convergence compared to the original DETR. This method improves on DETR’s slow convergence and low performance in detecting small objects, but still requires longer training schedules than CNN-based detectors, and remains computationally heavy.
- **RetinaNet [33]**. A one-stage object detector that uses focal loss to address class imbalance by focusing on hard examples, rather than the more plentiful easy negatives. It was the first one-stage detector to match the accuracy of more complex two-stage detectors. However, performance on very small objects remains lower than for large objects and does not out-perform some two-stage detectors.
- **YOLOv3 (YOLO) [34]**. One-stage object detector that uses multi-scale features for object detection, with each anchor responsible for a specific ground truth object. It is known for high speed and strong overall performance.
- **HRNet [35]**. High-resolution network that maintains multi-scale feature representations for keypoint prediction. It is a computationally efficient model that was originally designed for human pose estimation.
- **DeepPose [36]**. The first method to apply deep neural networks to human pose estimation by using cascaded regression. This improves pose reasoning and also captures contextual information.
- **DeepLabCut (DLC) [13]**. A commonly used pose estimation toolkit and model that is used in animal tracking. It is optimized for inference speed and includes a built-in GUI for labeling, training, and inference. This toolkit uses custom annotation format.
- **SLEAP [14]**. Another standard model used in neuroscience for animal tracking. It is optimized for fast inference and flexible workflows, includes a GUI for iterative labeling, training, and inference, and uses a custom annotation format.

We selected models that are practical for most research labs to adopt and are computationally efficient. Our selection also aimed to cover a breadth of implementations using lighter models, transformer-based detectors, and both top-down and bottom-up approaches. We configured each model using standard MMPose configuration files. We applied consistent training parameters where applicable and used data augmentations such as random rotation, scaling, and horizontal flipping to improve generalization. All training ran on NVIDIA V100 or A100 GPUs. Configuration files are provided in the GitHub repository.

### 8.5 Inference and Evaluation

We performed inference on the same systems used for training. For top-down models, we used object detectors to generate bounding boxes and passed cropped regions to pose estimators. We recorded predictions in JSON format and converted results into data frames for analysis.

To evaluate accuracy, we computed the Proportion of Correct Keypoints (PCK). In order to evaluate if a keypoint prediction is correct, we normalize by distance from the ground truth keypoint based on the body length calculated from the ground truth annotations. We defined a keypoint prediction as correct if it fell within a threshold distance (relative to body length) from the ground truth. We reported PCK at thresholds of 0.1 to 1.0 body length (step 0.1), and focus on 0.5 (half a body length) for main comparisons. This range visualizes performance at varying thresholds, depending on the level of accuracy that is important to the user. Depending on the task, the important PCK threshold may vary.

To evaluate inference speed, we measured frames per second (FPS) on an NVIDIA V100 GPU. We report FPS with PCK to show trade-offs between accuracy and throughput relevant for large-scale inference.

#### 8.5.1 Mapping Keypoints from TopViewMouse-5K to Maze and OFA

The three datasets we train on have independent keypoint sets that were selected for each task. To create the combined datasets, we map the keypoints from the OFA and Maze datasets to match the TopViewMouse-5K keypoint set (Table 1). Not all keypoints have a one-to-one comparison; we select the keypoint location that is the closest to the one present in the TopViewMouse-5K (Table 4).

### 8.6 Software and Library Environment

Experiments used MMPose version 1.3.2 from the OpenMMLab project as the foundation for our experiments[18]. We constructed a containerized environment using Docker/Singularity that included MMPose, MMDetection, pycocotools, and other necessary libraries. This setup ensures reproducibility and simplified deployment across systems. Environmental files and configuration templates are provided in the GitHub repository.

## Notes

### Competing Interest Statement

The authors have declared no competing interest.

